## Supplementary Material for "Is the brain macroscopically linear? A system identification of resting state dynamics"

### Supplementary Note 1: Effect of Bandpass Filtering on System Identification and the Detection of Nonlinearity

A standard step in the preprocessing of resting state fMRI time series is bandpass filtering, typically over the range $[0.01, 0.08]$ Hz [1], to reduce the contribution of non-neuronal sources on the signal and improve the SNR. In this work, however, we purposefully avoided this step. In the following, we discuss in detail the role and effects of pre-filtering in this rather unconventional context of system identification and, in particular, detection of nonlinear dynamics.

1. First, bandpass filtering (or any linear filtering for this matter) has no effect on the fitting or evaluation of linear models. The reason, in short, is the commuting property of linear systems. More specifically, any linear system including all of the ones used in this work can be written in the impulse-response form [2]

$$\mathbf{y}(t) = \mathbf{G}(q)\mathbf{e}(t),$$

where $\mathbf{G}$ is the transfer matrix

$$\mathbf{G}(q) = \begin{bmatrix} G_{11}(q) & G_{12}(q) & \cdots \\ G_{21}(q) & G_{22}(q) & \cdots \\ \vdots & \vdots & \ddots \end{bmatrix}$$

such that for any $i$ and $j$

$$G_{ij}(q) = g_{ij}(0) + g_{ij}(1)q^{-1} + g_{ij}(2)q^{-2} + \cdots$$

Here, $g_{ij}(t)$ is the impulse response from the the $j$th input $e_j(t)$ to the $i$th output $y_i(t)$, and $q$ is the standard shift operator such that $q^{-1}s(t) = s(t-1)$ for any signal $s(t)$ [3]. Recall that $\mathbf{y}(t)$ is the BOLD time series without bandpass filtering, as used in the main text, and let

$$F(q) = f(0) + f(1)q^{-1} + f(2)q^{-2} + \cdots$$

be any linear filter, including the bandpass filter used in common preprocessing pipelines. Assume, without loss of generality, that $f(0) \neq 0$ (an almost identical argument can be given if $f(0)$ or any number of the first terms in $F(q)$ are zero, simply by factoring out enough powers of $q^{-1}$). The output of the filter is

$$\mathbf{z}(t) = F(q)\mathbf{y}(t) = F(q)\mathbf{G}(q)\mathbf{e}(t).$$

It then immediately follows from the prediction error framework [3] that the one step ahead prediction error of $\mathbf{z}(t)$ is given by

$$\begin{aligned} \mathbf{z}(t) - \hat{\mathbf{z}}(t|t-1) &= f(0)\mathbf{G}^{-1}(q)F^{-1}(q)\mathbf{z}(t) \\ &= f(0)\mathbf{G}^{-1}(q)\mathbf{y}(t) \\ &= f(0)\big[\mathbf{y}(t) - \hat{\mathbf{y}}(t|t-1)\big]. \end{aligned}$$

In other words, the prediction error of $\mathbf{z}(t)$ is identical to the prediction error of $\mathbf{y}(t)$, except for a constant factor equal to the instantaneous gain of the filter. Therefore, not only is fitting a model by minimizing the prediction error of $\mathbf{z}(t)$ identical to fitting a model by minimizing the prediction error of $\mathbf{y}(t)$, but the cross-validated $R^2$ (up to a fixed constant) and residual whiteness of these models are also identical.

This argument clearly fails for nonlinear systems. Indeed, fitting a nonlinear model on $\mathbf{z}(t)$ can result in a different model with different $R^2$ and residual whiteness than the same model fit on $\mathbf{y}(t)$. The critical point, however, is that the linear filter $F(q)$ cannot generate nonlinearity, while it can certainly weaken and even eliminate it. We explain these two points in more detail next.

2. Assume, first, that the dynamics of $\mathbf{y}(t)$ are truly linear, as seems to be the case from our analysis in the main text. Then, the relationship between $\Delta\mathbf{y}(t)$ and $\mathbf{y}(t-1)$ (as random vectors) is linear. By definition, if the relationship between two random vectors $\mathbf{u}$ and $\mathbf{v}$ is linear, they can be written in the form

$$\begin{bmatrix}\mathbf{u}\\ \mathbf{v}\end{bmatrix} = \begin{bmatrix}\mathbf{A}_{11} & \mathbf{A}_{12}\\ \mathbf{A}_{21} & \mathbf{A}_{22}\end{bmatrix}\begin{bmatrix}\mathbf{e}_1\\ \mathbf{e}_2\end{bmatrix} \tag{S1}$$

where $\mathbf{A}$ is an appropriate matrix and $\mathbf{e}_1$ and $\mathbf{e}_2$ are independent. Now let $\mathbf{u}_F$ be the result of applying a linear filter $F(q)$ to (the samples of) $\mathbf{u}$. In other words, if $N$ is the order of an (arbitrarily accurate) FIR approximation of $F(q)$,

$$\mathbf{u}_F = \begin{bmatrix}\mathbf{u}_1 & \mathbf{u}_2 & \cdots & \mathbf{u}_N\end{bmatrix}\begin{bmatrix}f(0)\\ f(1)\\ \vdots\\ f(N-1)\end{bmatrix} = \begin{bmatrix}\mathbf{u}_1 & \mathbf{u}_2 & \cdots & \mathbf{u}_N\end{bmatrix}\mathbf{f} \tag{S2}$$

where $f(t)$ is the impulse response of $F(q)$ and $\mathbf{u}_1, \mathbf{u}_2, \dots, \mathbf{u}_N$ are identically distributed (but not necessarily independent) samples of $\mathbf{u}$. From Eq. (S1),

$$\begin{bmatrix}\mathbf{u}_1 & \mathbf{u}_2 & \cdots & \mathbf{u}_N\end{bmatrix} = \begin{bmatrix}\mathbf{A}_{11} & \mathbf{A}_{12}\end{bmatrix}\begin{bmatrix}\mathbf{e}_{1,1} & \mathbf{e}_{1,2} & \cdots & \mathbf{e}_{1,N}\\ \mathbf{e}_{2,1} & \mathbf{e}_{2,2} & \cdots & \mathbf{e}_{2,N}\end{bmatrix} \tag{S3}$$

where $\mathbf{e}_{1,1}, \dots, \mathbf{e}_{1,N}$ are identically distributed (but not necessarily independent) samples of $\mathbf{e}_1$, similarly for $\mathbf{e}_2$. Note that each $\mathbf{e}_{1,t}$ is still independent from each $\mathbf{e}_{2,s}$ by definition. Substituting Eq. (S3) into Eq. (S2) thus gives

$$\begin{aligned}\mathbf{u}_F &= \begin{bmatrix}\mathbf{A}_{11} & \mathbf{A}_{12}\end{bmatrix}\begin{bmatrix}\mathbf{e}_{1,1} & \mathbf{e}_{1,2} & \cdots & \mathbf{e}_{1,N}\\ \mathbf{e}_{2,1} & \mathbf{e}_{2,2} & \cdots & \mathbf{e}_{2,N}\end{bmatrix}\mathbf{f}\\ &= \begin{bmatrix}\mathbf{A}_{11} & \mathbf{A}_{12}\end{bmatrix}\begin{bmatrix}\mathbf{e}_{1,F}\\ \mathbf{e}_{2,F}\end{bmatrix}\end{aligned}$$

where $\mathbf{e}_{1,F}$ and $\mathbf{e}_{2,F}$ are filtered versions of $\mathbf{e}_1$ and $\mathbf{e}_2$ and, still, independent of each other. Following the same steps for $\mathbf{v}$, we get

$$\begin{bmatrix}\mathbf{u}_F\\ \mathbf{v}_F\end{bmatrix} = \begin{bmatrix}\mathbf{A}_{11} & \mathbf{A}_{12}\\ \mathbf{A}_{21} & \mathbf{A}_{22}\end{bmatrix}\begin{bmatrix}\mathbf{e}_{1,F}\\ \mathbf{e}_{2,F}\end{bmatrix}$$

showing that $\mathbf{u}_F$ and $\mathbf{v}_F$ are still linearly related after filtering. This is indeed expected since a linear filter cannot generate nonlinear dependence between two signals that are originally linearly related.

3. Now assume, in contrast, that the dynamics of $\mathbf{y}(t)$ is in fact nonlinear and we filter $\mathbf{y}(t)$ to get $\mathbf{z}(t) = F(q)\mathbf{y}(t)$. The best that can happen, as far as detecting nonlinearity is concerned, is that the dynamics of $\mathbf{z}(t)$ remain nonlinear. However, it is possible that $F(q)$ weakens or completely averages out the nonlinearities in $\mathbf{y}(t)$, as we saw in the main text. In fact, the common bandpass filter over $[0.01, 0.08]$ Hz (compared to a Nyquist frequency of $1/2\text{TR} \simeq 0.7$ Hz) is strongly lowpass and involves significant averaging over time.

In conclusion, while bandpass filtering has no effect on linear models and preserves linearity of time series, it can well weaken/eliminate any nonlinearity in the time series. Therefore, regardless of how much "cleaner" bandpass-filtered data might be, finding no nonlinearity before bandpass filtering, as pursued in the main text, is a stronger statement than the same finding would be if obtained after bandpass filtering.

### Supplementary Figures

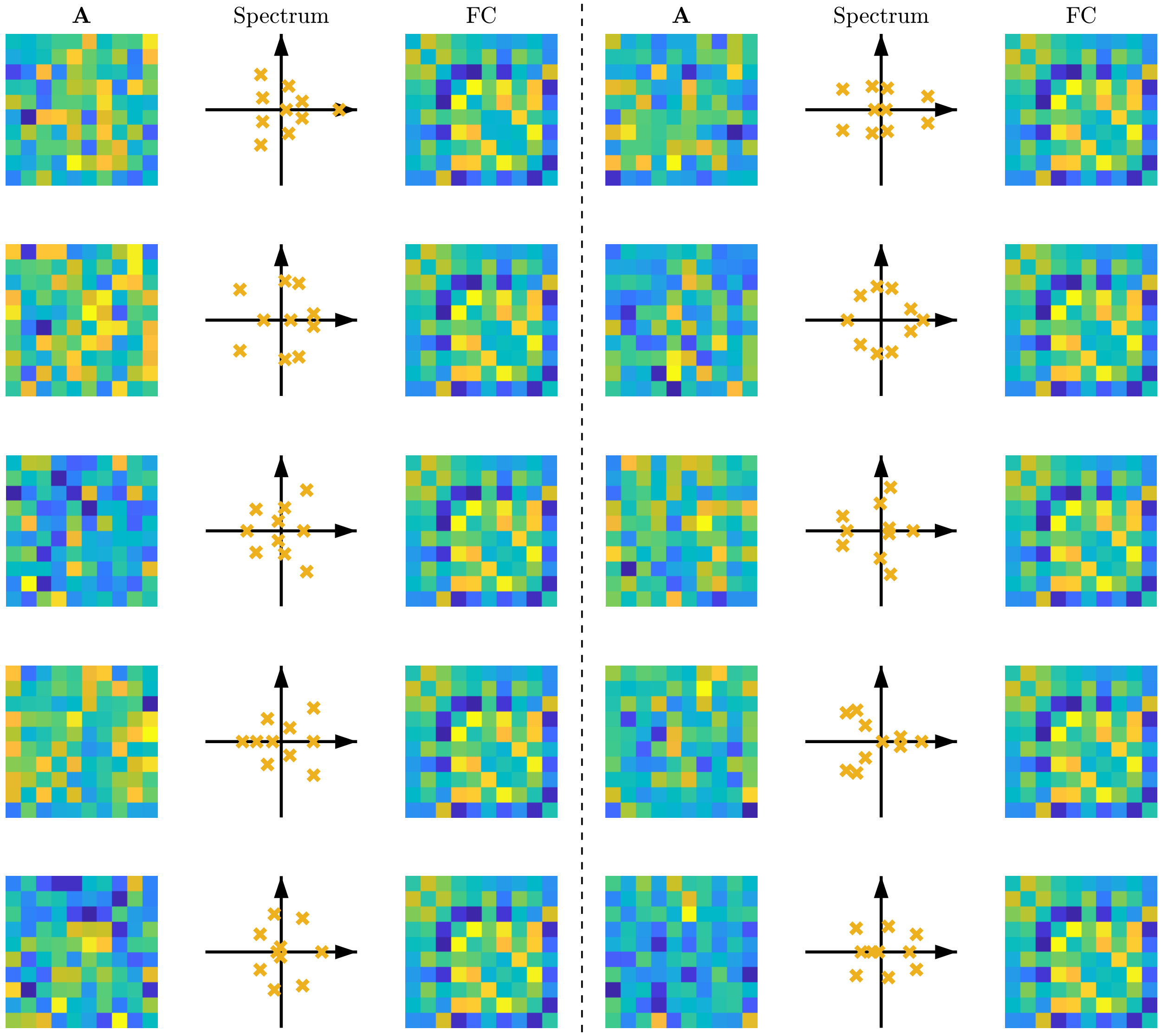


**Supplementary Figure 1: Different systems can produce almost identical functional connectivity (FC).** Ten different systems of the form $\mathbf{y}(t) = \mathbf{A}\mathbf{y}(t-1) + \mathbf{e}(t)$ are shown with completely different $\mathbf{A}$ matrices and spectra (eigenvalues of $\mathbf{A}$), but almost identical FC matrices. The Pearson correlation coefficient between all pairs of FCs is 1.0000 while the absolute value of the Pearson correlation coefficients between the $\mathbf{A}$ matrices range from a minimum of 0.0023 to a maximum of 0.36. All systems were driven by independent and randomly generated noise sequences $\mathbf{e}(t)$. In the figures, the diagonal entries of the FC matrices (which are by definition equal to 1) are set 0 to increase color resolution.

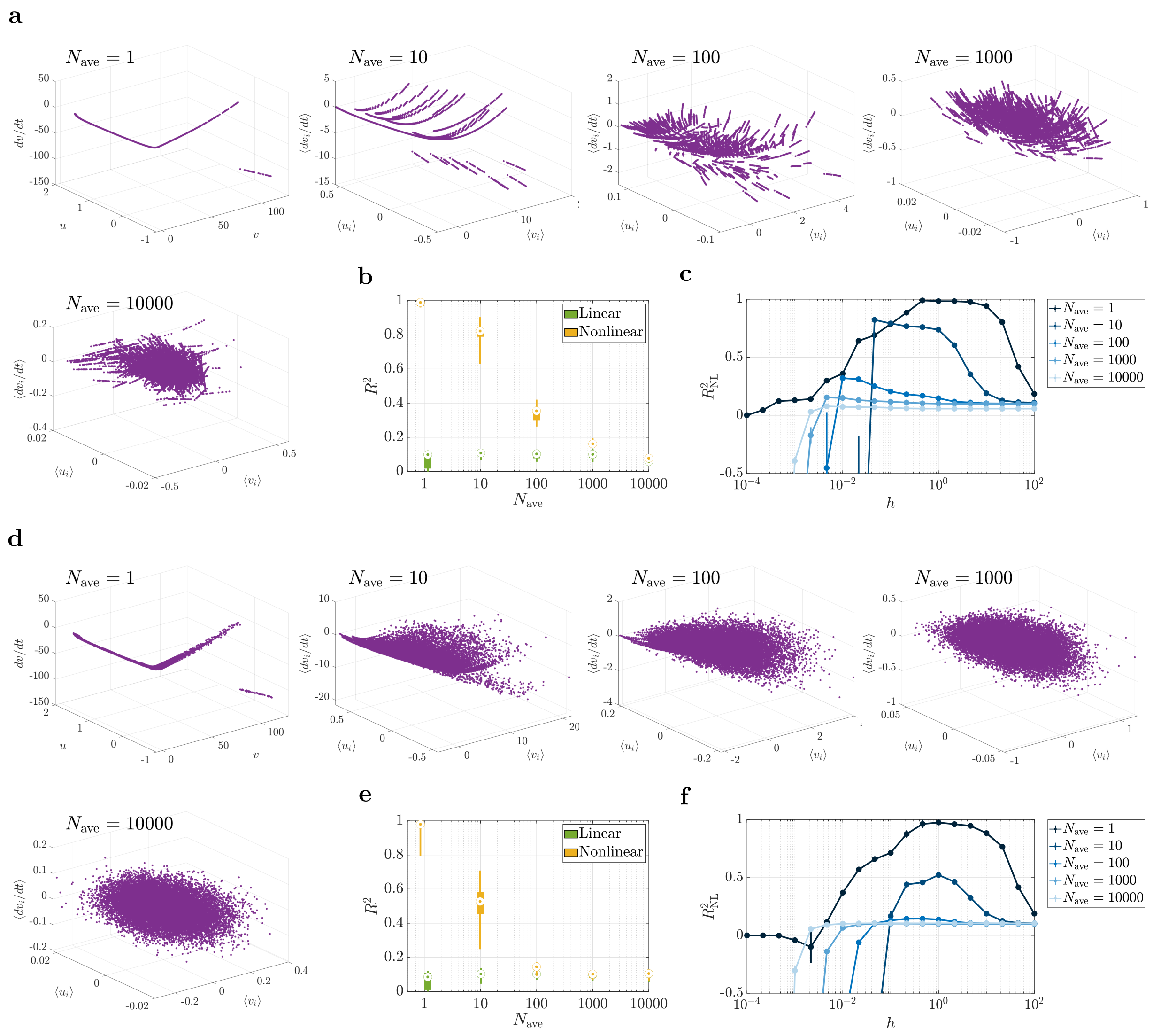


**Supplementary Figure 2: The linearizing effect of spatial averaging on the Izhikevic spiking model. (a)** The Izhikevic model [4] was simulated for $N_{\text{ave}}$ neurons and their $v_i$ and $u_i$ variables were averaged before being "observed". The neurons had the same parameters (and hence the same spiking frequency) but were initialized at random phases of their limit cycles. The average of $dv_i/dt$ is plotted against the average of $v_i$ and $u_i$, showing the loss of nonlinearity at about $N_{\text{ave}} \sim 10^3$-$10^4$. $dv_i/dt$ is chosen since it has a more nonlinear dependence on $v_i$ and $u_i$ than $du_i/dt$ for one neuron, but the same (even stronger) linearizing effect holds for $(\langle v_i \rangle, \langle u_i \rangle, \langle du_i/dt \rangle)$. **(b)** The $R^2$ distributions of a linear (simple linear regression) and nonlinear (manifold-based locally linear regression) model for the relationships in panel (a) and 100 random repetitions. The manifold-based model was chosen over the MMSE model as it consistently gave significantly higher $R^2$ values (due to the high sample complexity of the MMSE model). Note that the $R^2$ distribution of the linear model is mostly flat (less affected by spatial averaging), while that of the nonlinear model decays rapidly with $N_{\text{ave}}$ until it reaches the linear level. **(c)** Nonlinear $R^2$ as a function of the manifold method's window sizes $h$ for varying $N_{\text{ave}}$ values. The circles and error bars show median and inter-quartile ranges, respectively. While an optimal, mid-range $h$ exists for small values of $N_{\text{ave}}$, the curve plateaus for large $h$ as $N_{\text{ave}}$ increases, showing the loss of nonlinearity with increasing $N_{\text{ave}}$. **(d-f)** Same as panels (a-c) but for a model where *process* noise is added to the $dv_i/dt$ equations. The noise has a very low power (SNR = 100) and is only meant to destroy the artificial micro-nonlinear relationships that are visible in panel (a) and can inflate the power of nonlinear regression for $N_{\text{ave}} \sim 10$-$10^3$. Note that this is different from the addition of observation noise discussed in the main text, even though adding observation noise could serve this purpose as well. In all box plots, the center point, box limits, and whiskers represent the median, upper and lower quartiles, and the smallest and largest samples, respectively.

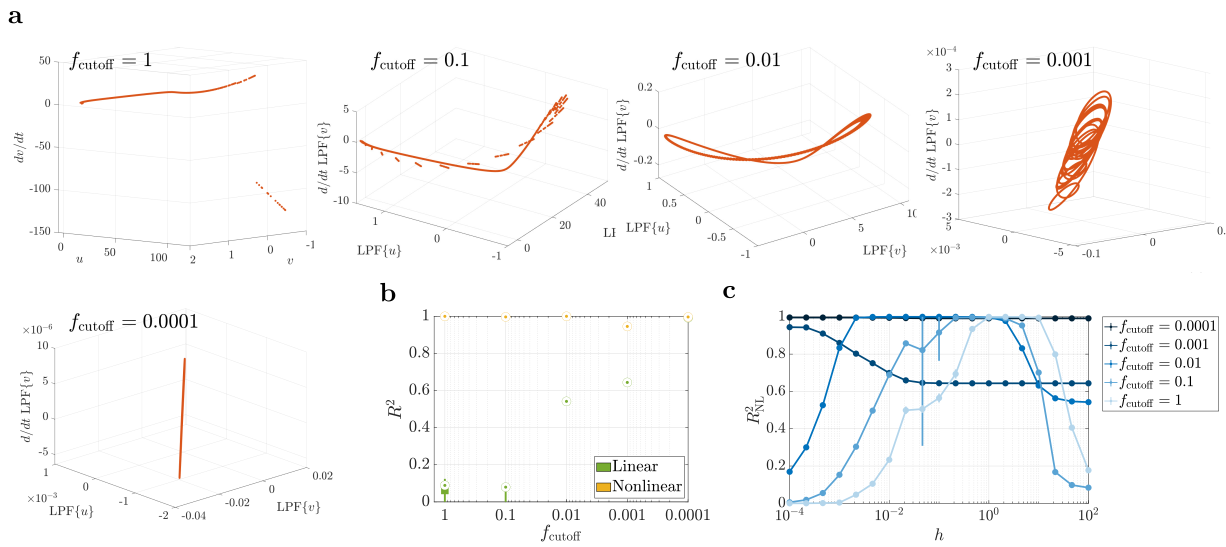


**Supplementary Figure 3: The linearizing effect of temporal averaging on the Izhikevic spiking model.** Panels parallel those in Figure 2. **(a)** One Izhikevic model [4] was simulated, starting from a random initial condition on its limit cycle, and its $v$ and $u$ variables were low-pass filtered before being "observed". $d/dt$ LPF$\{v\}$ is plotted against LPF$\{v\}$ and LPF$\{u\}$, showing the loss of nonlinearity at about $f_{\mathrm{cutoff}} \sim 10^{-4}$. **(b,c)** Similar to panels (b,c) in Figure 2, except that here the $R^2$ of the linear model reaches that of the nonlinear one as temporal averaging is intensified. The nonlinear model always has a near-perfect prediction power given the fully deterministic nature of this simulation (i.e., a sufficiently small $h$ was always sufficient to perfectly match a locally-linear model to the curves in panel (a). Note that addition of process noise here does not have the same "blurring" effect of Figure 2(d-f) since it lies before the low-pass filter. Observation noise can nevertheless be added with the same effect (not shown here). In all box plots, the center point, box limits, and whiskers represent the median, upper and lower quartiles, and the smallest and largest samples, respectively.

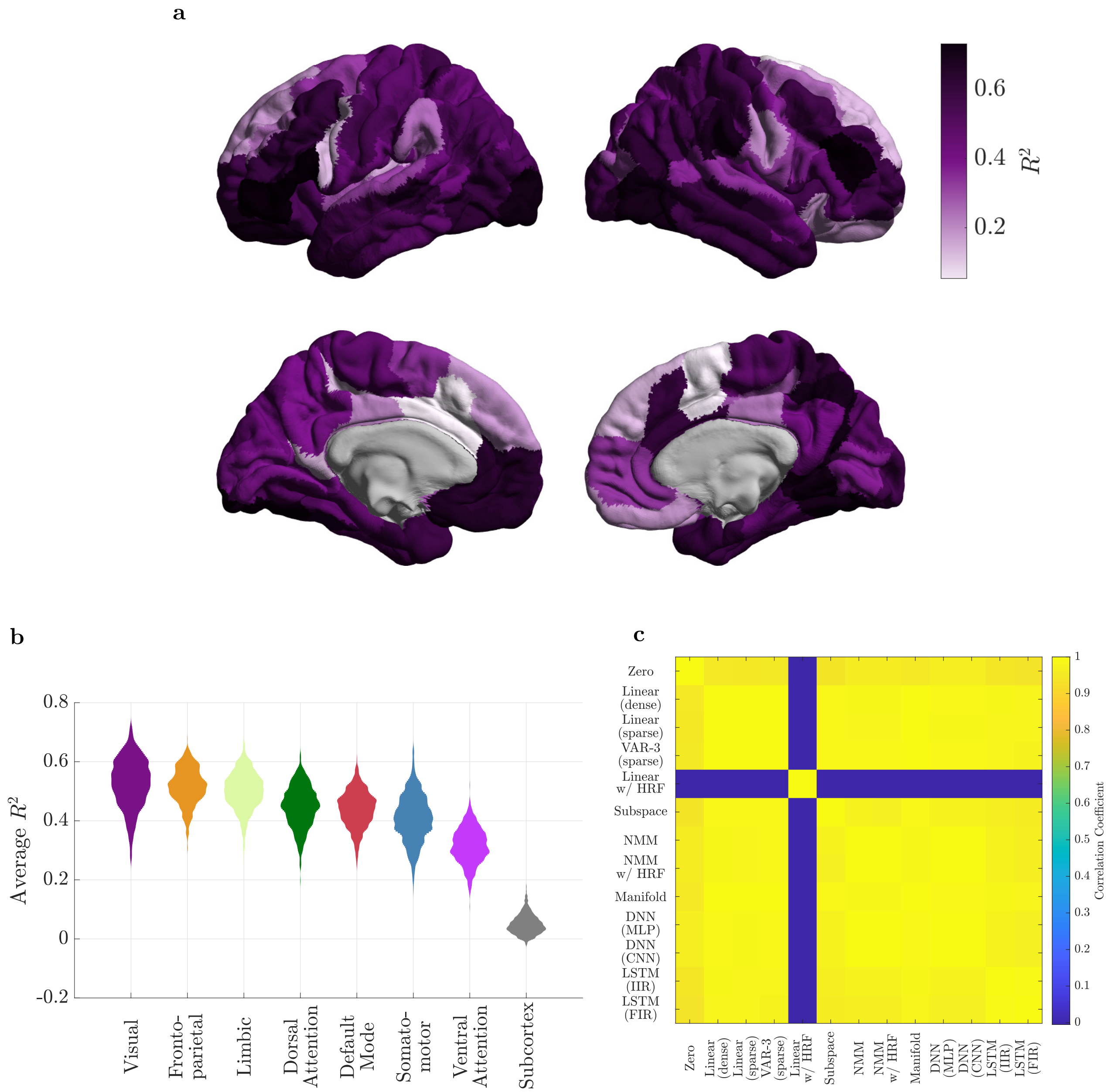


**Supplementary Figure 4: The spatial distribution of predictability.** **(a)** The cortical distribution of the $R^2$ of our best model ('VAR-3 (sparse)'), averaged over the 700 subjects. **(b)** Violin plots of the distribution of $R^2$, averaged over all the regions of each resting state network, for all subjects. Note that each distribution in panel **(b)** is thus composed of 700 samples. All pairs of distributions have significantly different medians in the order plotted (one-sided Mann-Whitney $U$-test at $\alpha = 0.05$ with BH-FDR correction for multiple comparisons). The most striking difference is between the cortical and subcortical regions, where the dynamics of the latter are remarkably less predictable than the former. This lower predictability can be viewed as having "more noisy" dynamics or, more precisely, less spatially and temporally correlated (i.e., more white) fMRI time series in these regions. **(c)** The correlation coefficient between the $R^2$ values of all the (brain-wise) methods. All methods produce almost the same cortical distributions of $R^2$ (albeit with different absolute values of $R^2$, cf. Figure 2a in the main text), except for 'Linear w/ HRF' which has an un-correlated $R^2$ distribution relative to the rest of the methods.

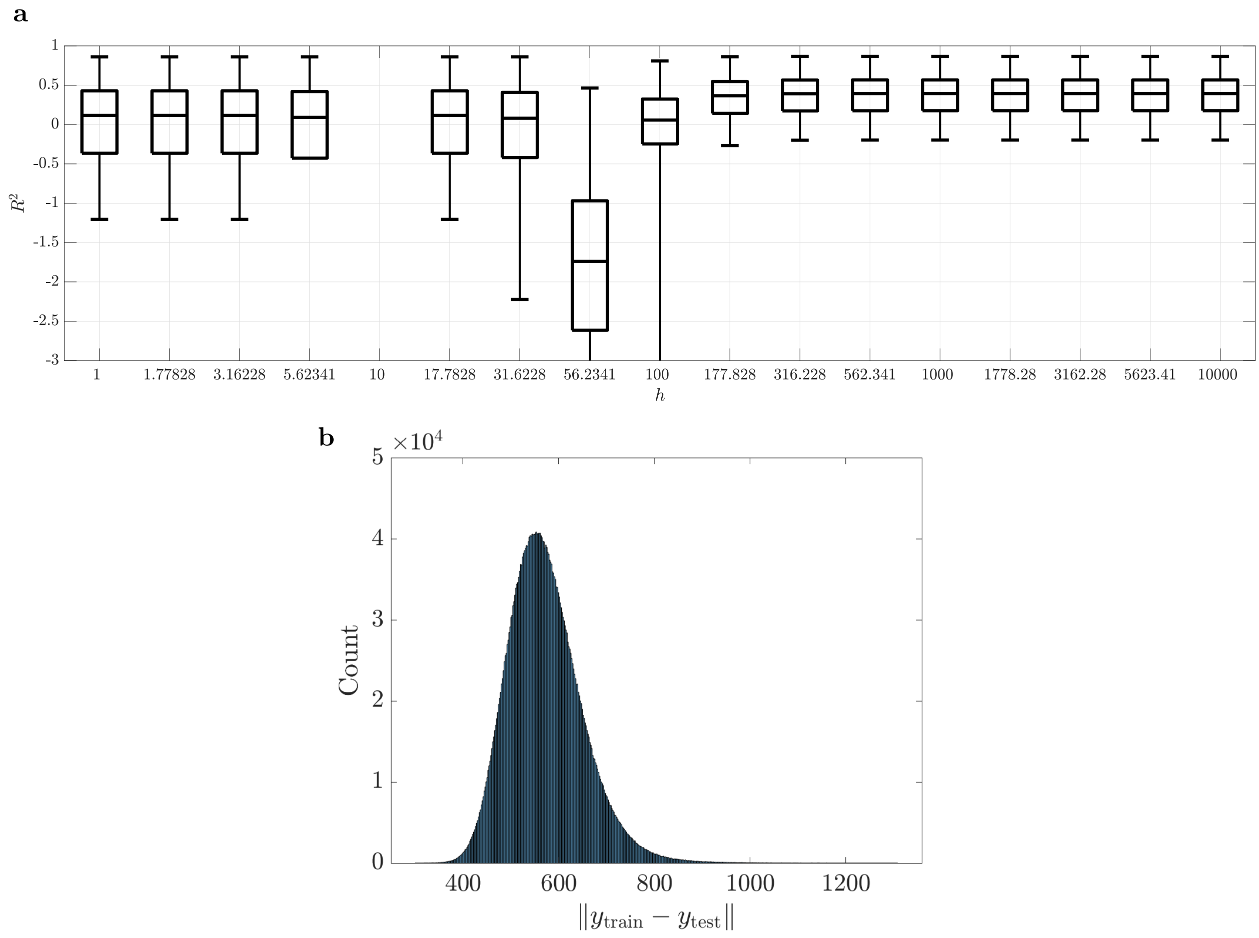


**Supplementary Figure 5: Effect of window size $h$ on the accuracy of the manifold-based locally linear ('Manifold') model in fMRI data.** (a) Boxplot of the $R^2$ distribution as a function of $h$, combined across the 116 brain regions of 70 randomly selected subjects (10% of all subjects to reduce computational cost). The $R^2$ values were not computable ('NaN') for $h = 10$ due to limited machine precision. The model is equivalent to the zero model for the three leftmost boxplots as no training point falls within the Gaussian-weighted neighborhood of any test points. As $h$ is increased to $h \sim 10$, few training data points start to fall within the neighborhood window of some of the test points, but are far enough that their Gaussian weights fall below machine precision, leading to missing ('NaN') predicted values and, hence, $R^2$. As $h$ is further increased, more training points fall within the neighborhood of each test point, but are few enough to lead to poor $R^2$, until $h$ is increased enough to reach the globally linear regime. (c) The distribution of the Euclidean distance between all pairs of training and test points for a randomly selected subject (subject 103818) to aid in understanding the trends that are apparent in panels (a) and (b). In particular, note that $h = 10^4$ (and even smaller values) clearly lead to a globally linear model as almost all of the training-test pairs of points have distances less than $h/10$.

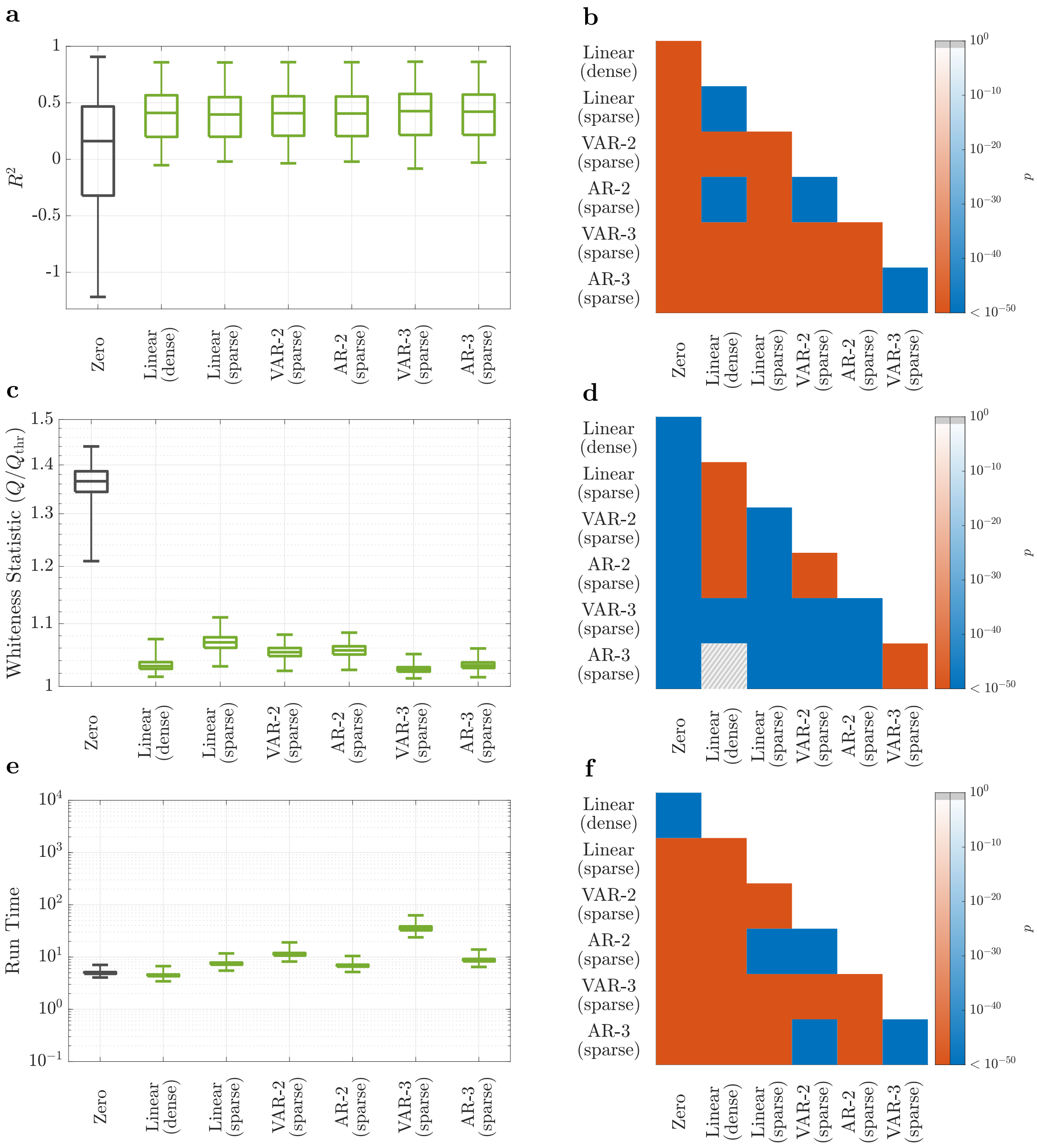


**Supplementary Figure 6: The effects of the number of lags and sparsity patterns on the prediction accuracy and computational complexity of linear AR models of rsfMRI.** Panels parallel those in Figure 2 in the main text and the descriptions of method acronyms are given in Table 1 therein. Generally, the number of lags and sparsity patterns have little effect on the prediction accuracy of linear AR models for rsfMRI data (in contrast to rsiEEG data, as explained in the main text), as seen from panel (a). The estimates of statistical significance in panel (b) are to a great extent due to the large sample size ($700 \times 116$). However, increasing the number of regressors, both by increasing the number of AR lags and by allowing for off-diagonal entries of all lags ('VAR' models), does lead to a non-trivial improvement in the whiteness of residuals, even though is often accompanied by non-trivial increases in model complexity and computation time as well. In all box plots, the center line, box limits, and whiskers represent the median, upper and lower quartiles, and the smallest and largest samples, respectively, and sample size = 81200.

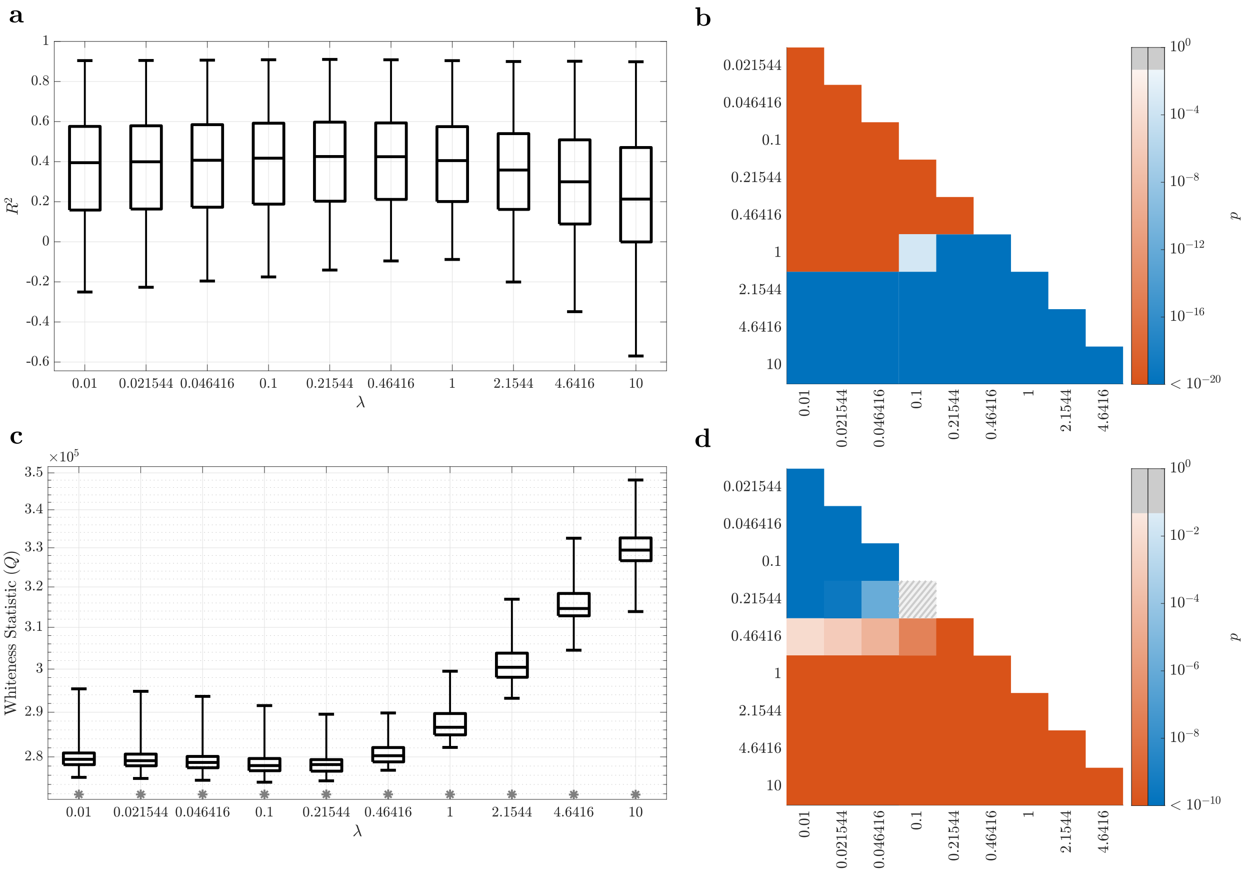


**Supplementary Figure 7: The effect of the LASSO parameter $\lambda$ on the accuracy of the 'VAR-3 (sparse)' model.** Panels parallel those in Figure 2 of the main text. **(a)** The distribution of the cross-validated regional $R_i^2$, combined across all regions and 10% of subjects (randomly selected), for varying values of $\lambda$. **(b)** The $p$-values of the one-sided Wilcoxon signed rank test performed between all pairs of distributions of $R^2$ in panel **(a)**. **(c, d)** Similar to panels **(a, b)** but for the $Q$ statistic of the test of whiteness of the residuals.

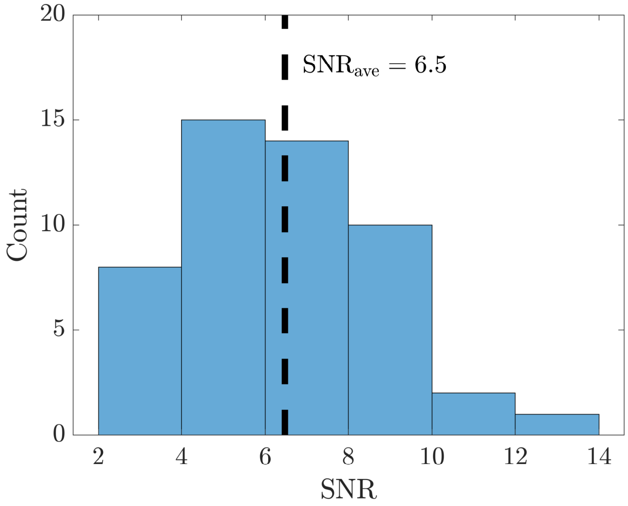


**Supplementary Figure 8: Histogram of scanner SNR estimates for rsfMRI data.** Scanner SNR was estimated for 50 randomly selected rest scans by comparing the average signal powers inside the respective subject's gray matter and outside of their head (see Methods). Due to the conservatism of this method, the resulting SNR estimates are expectedly over-estimated, but yet are not far from the SNR = 1 level that is enough to completely mask nonlinear interactions on its own (Figure 4g-h).

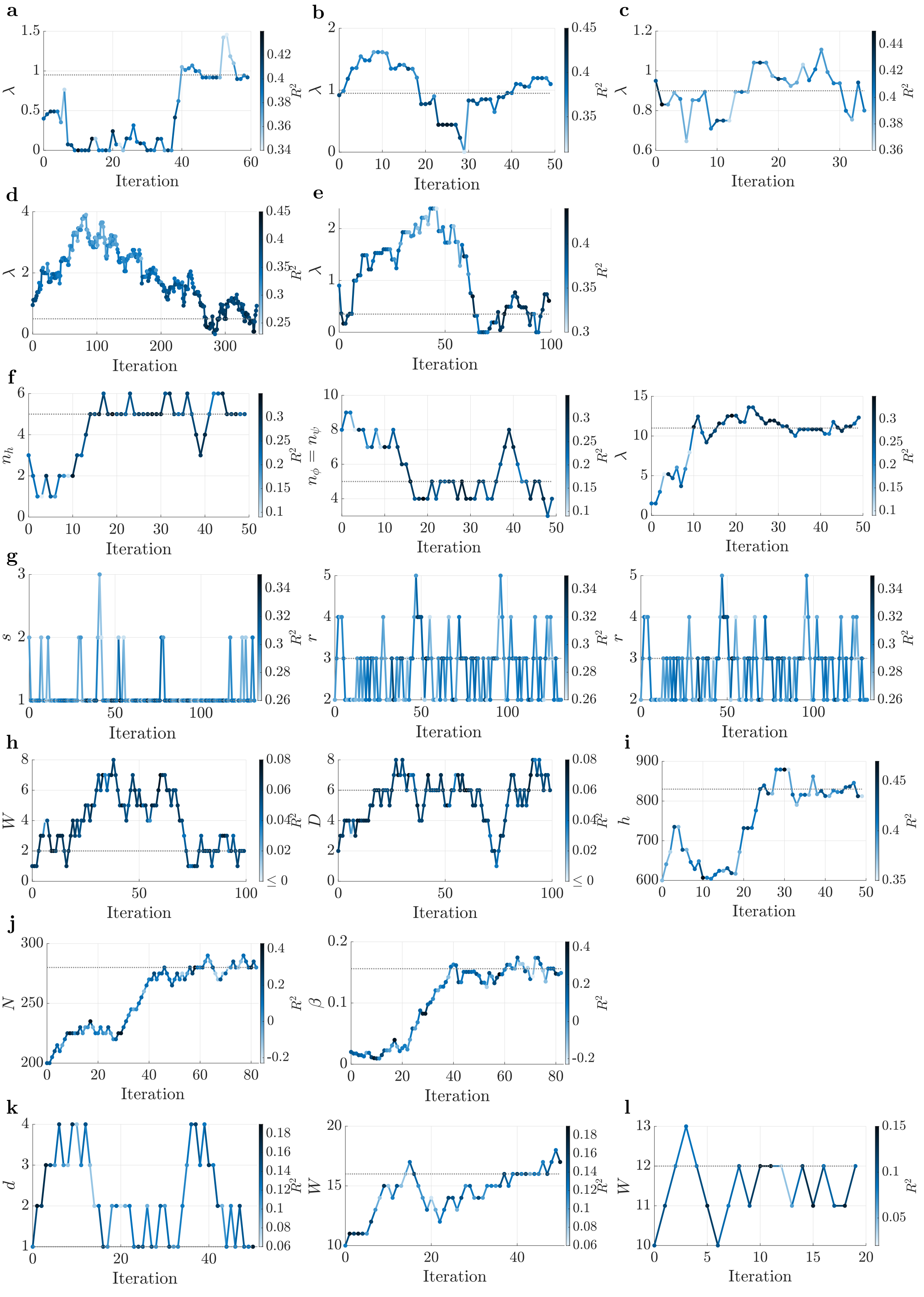
a
b
c
d
e
f
g
h
i
j
k
l
Iteration
$\lambda$
$R^2$
$n_h$
$n_\phi = n_\psi$
$s$
$r$
$W$
$D$
$h$
$N$
$\beta$
$d$
$\leq 0$

**m**

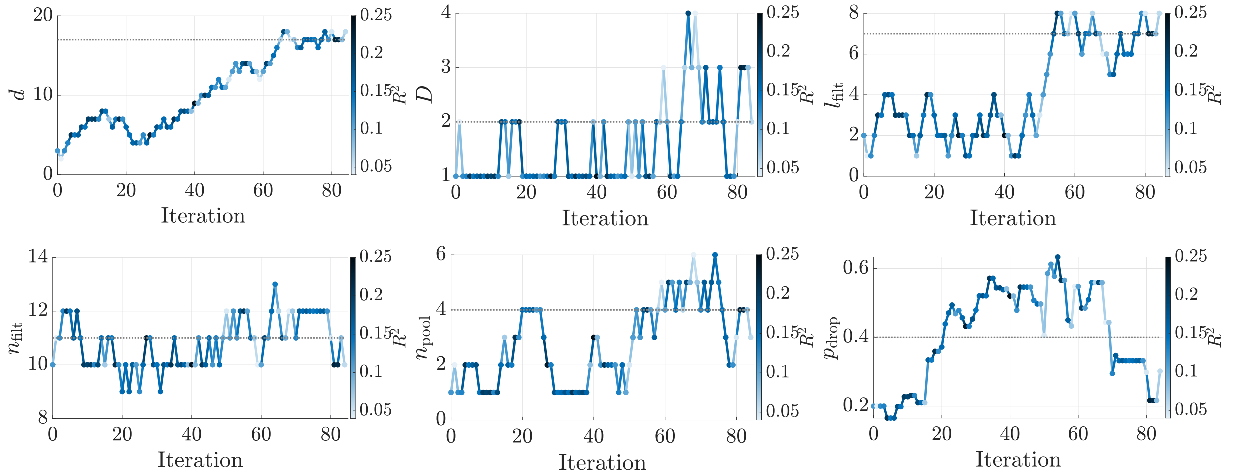


**Supplementary Figure 9: Hyper-parameter tuning of linear and nonlinear model families for fMRI.** For each parametric family of models, its hyper-parameters were simultaneously optimized using stochastic gradient descent (SGD, see Methods) to select the model with the highest cross-validated $R^2$ within that model family. (a) Linear (sparse); (b) AR-2 (sparse); (c) VAR-2 (sparse); (d) AR-3 (sparse); (e) VAR-3 (sparse); (f) Linear w/ HRF; (g) Subspace; (h) DNN (MLP); (i) Manifold; (j) MMSE (pairwise); (k) LSTM (FIR); (l) LSTM (IIR); (m) DNN (CNN). Each panel shows the evolution of the hyper-parameter(s) of one model family during the SGD iterations, color-coded with the value of $R^2$ at each iteration, and the hyper-parameter value(s) selected as optimal (dotted gray lines, also given in Table 1 in the main text). Note that the hyper-parameter(s) are not expected to converge to a fixed (optimal) value, but rather to fluctuate around it due to the stochastic nature of SGD and the natural variability of data segments.

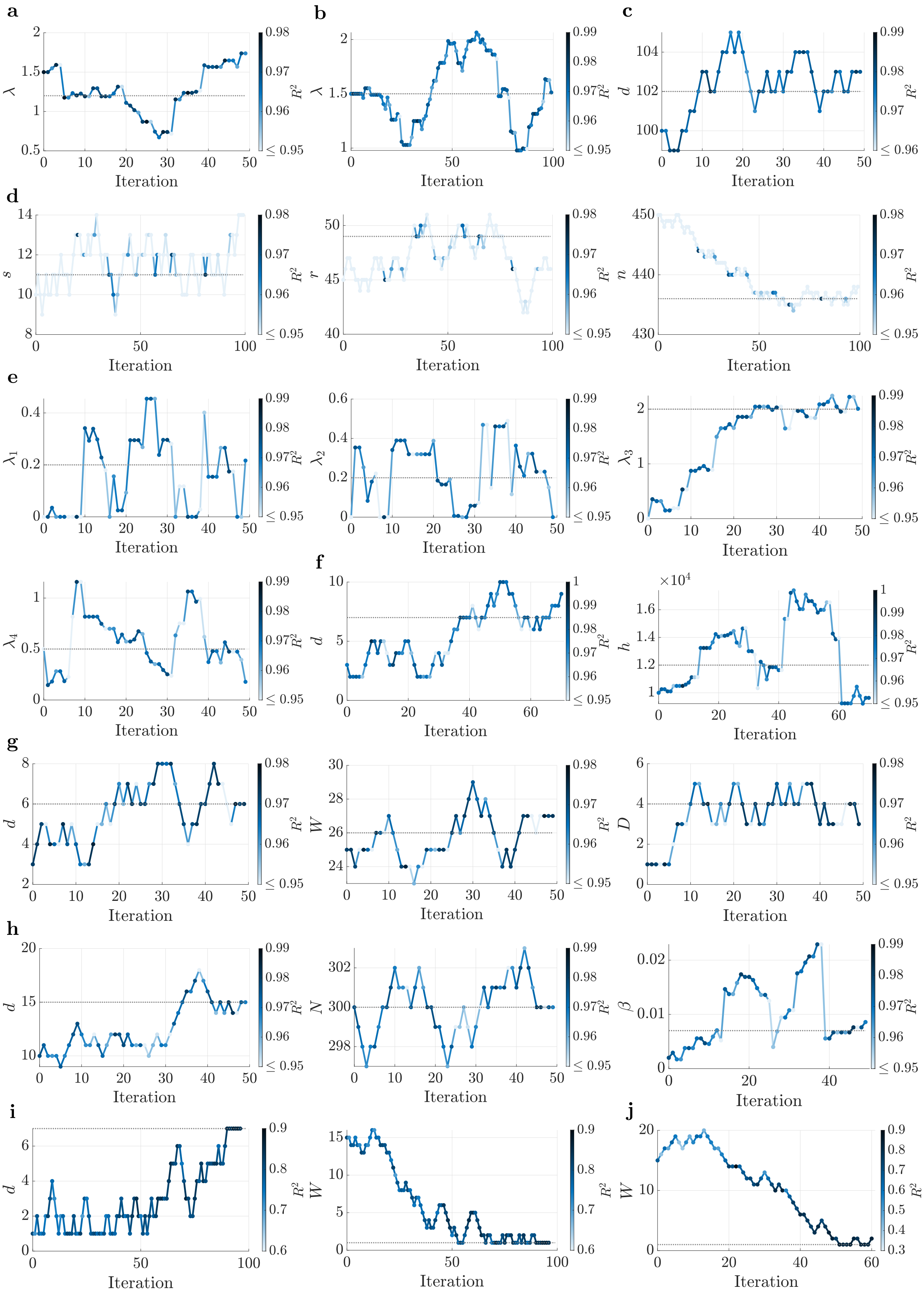
a
b
c
d
e
f
g
h
i
j
Iteration
$\lambda$
$d$
$s$
$r$
$n$
$\lambda_1$
$\lambda_2$
$\lambda_3$
$\lambda_4$
$h$
$W$
$D$
$N$
$\beta$
$R^2$
$\times 10^4$

**k**

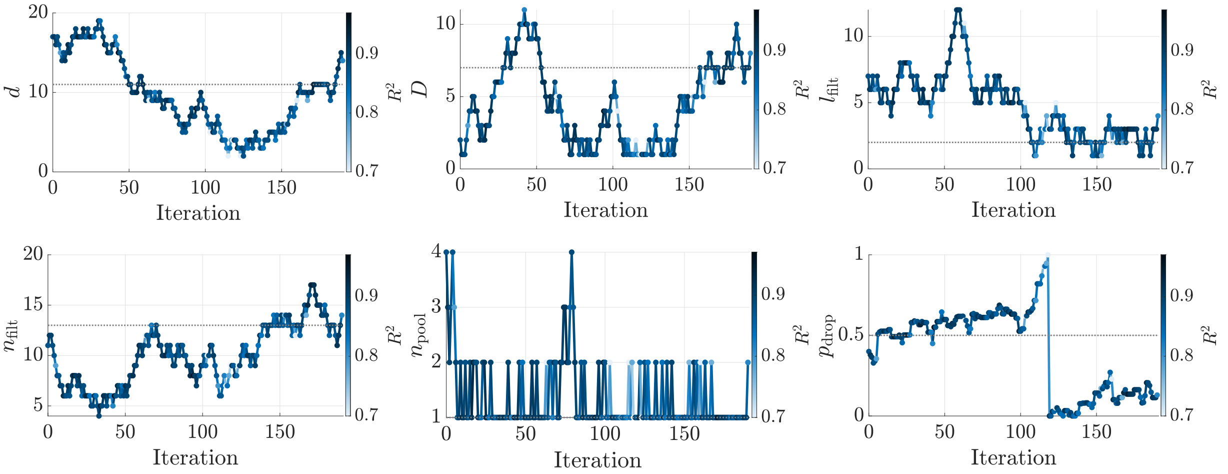


**Supplementary Figure 10: Hyper-parameter tuning of linear and nonlinear model families for iEEG.** (a) Linear (sparse); (b) AR-100 (sparse); (c) AR-100 (scalar); (d) Subspace; (e) NMM; (f) Manifold; (g) DNN; (h) MMSE (scalar); (i) LSTM (FIR); (j) LSTM (IIR); (k) DNN (CNN). Details and interpretations are the same as Supplementary Figure 9.

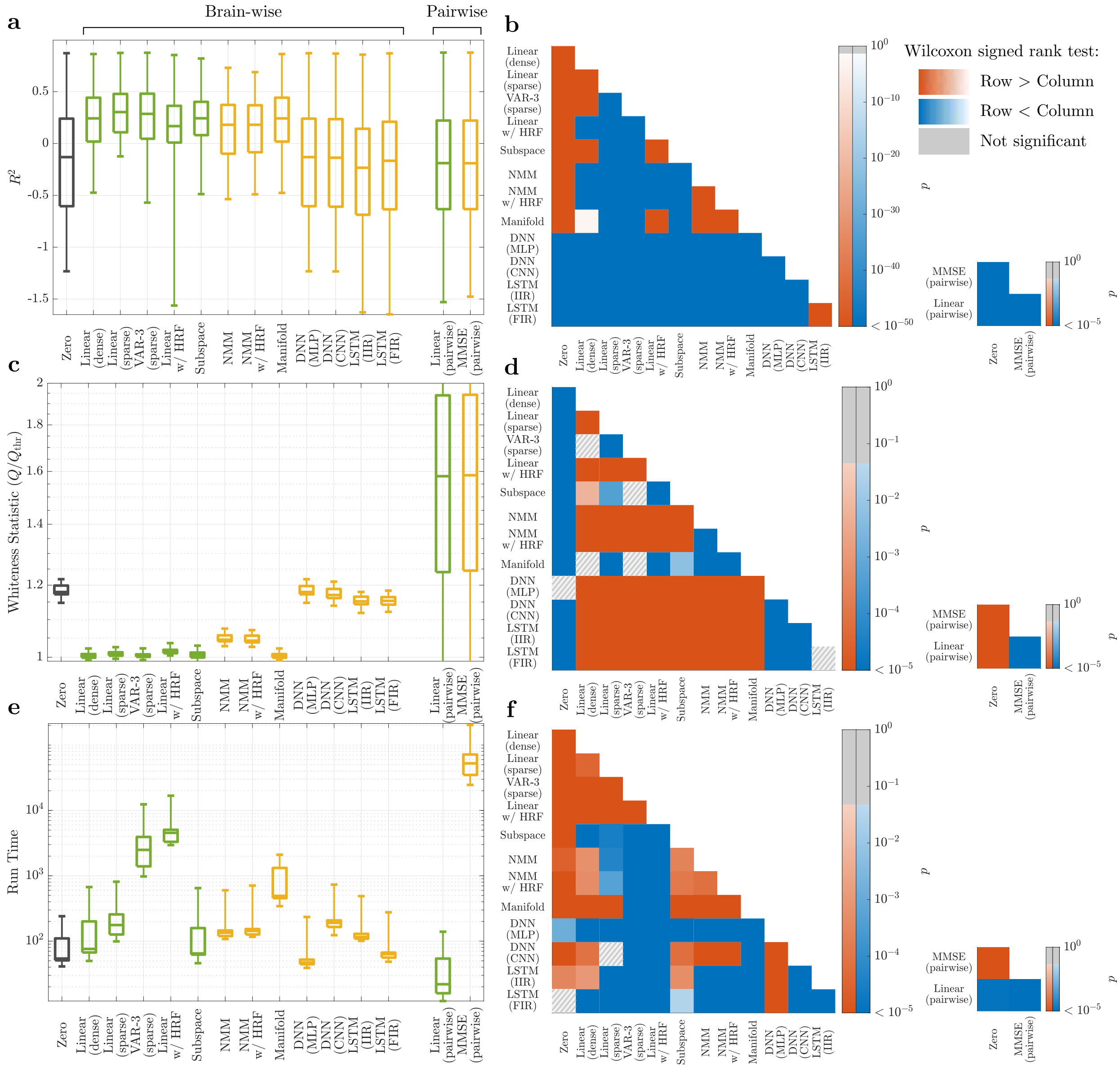

**Supplementary Figure 11: Linear vs. nonlinear models of finely-parcellated rsfMRI activity.** 400 cortical parcels (Schaefer 400x17 [5]) and 50 subcortical ones (Melbourne Scale III [6]) were used. Panels and details parallel those in Figure 2 in the main text, except that only data from 32 randomly selected subjects and a single-fold cross-validation has been used to reduce computational complexity. The half-session used as the test for each subject has been selected at random and the remaining 7 half-sessions have been used for training, as in the main text. The model with the highest $R^2$ is now the 'Linear (sparse)' even though 'VAR-3 (sparse)' still has whiter residuals.

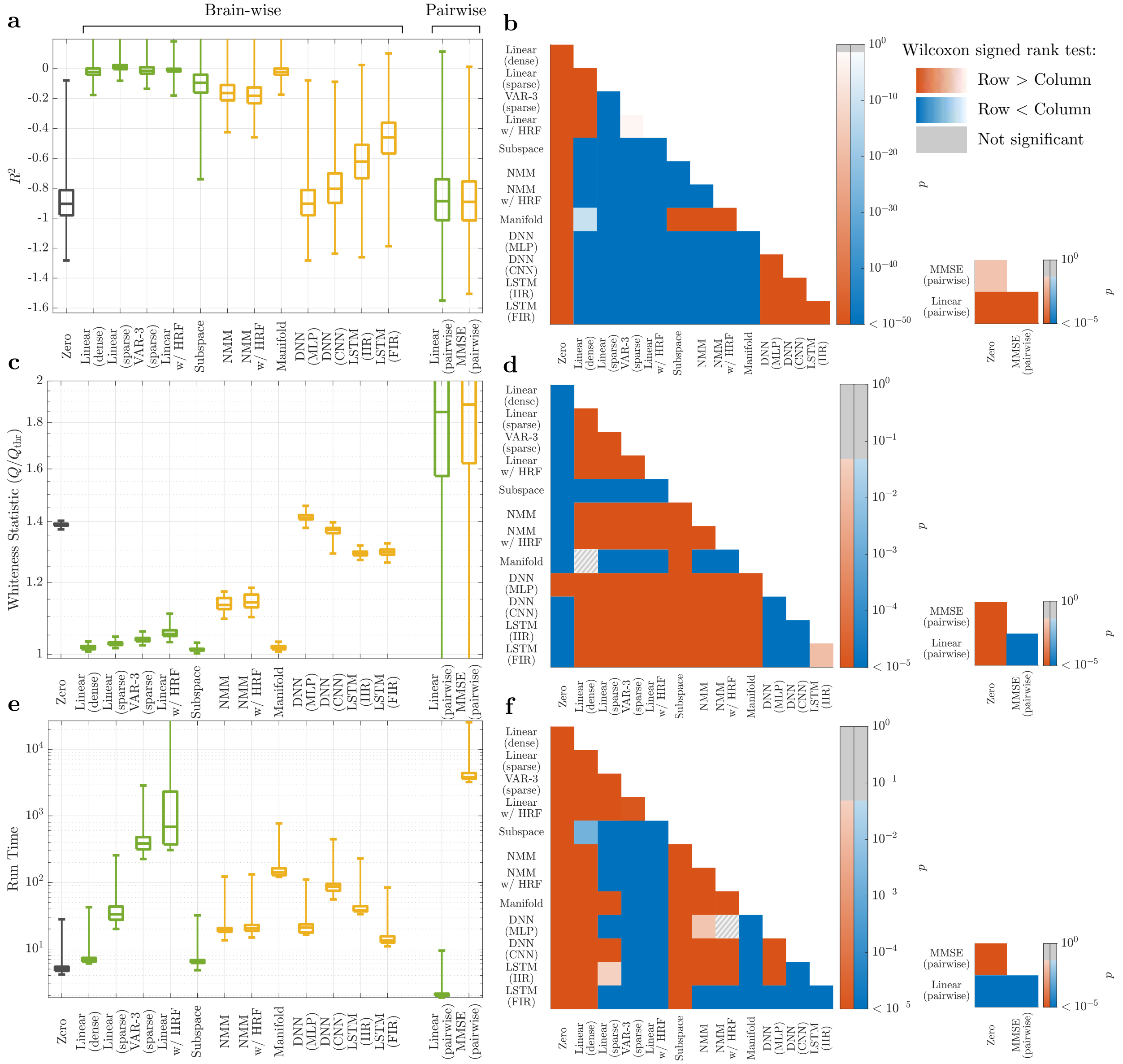
a
Brain-wise
Pairwise
$R^2$
b
Wilcoxon signed rank test:
Row > Column
Row < Column
Not significant
c
Whiteness Statistic ($Q/Q_{\text{thr}}$)
d
e
Run Time
f
Zero
Linear (dense)
Linear (sparse)
VAR-3 (sparse)
Linear w/ HRF
Subspace
NMM
NMM w/ HRF
Manifold
DNN (MLP)
DNN (CNN)
LSTM (IIR)
LSTM (FIR)
Linear (pairwise)
MMSE (pairwise)
p

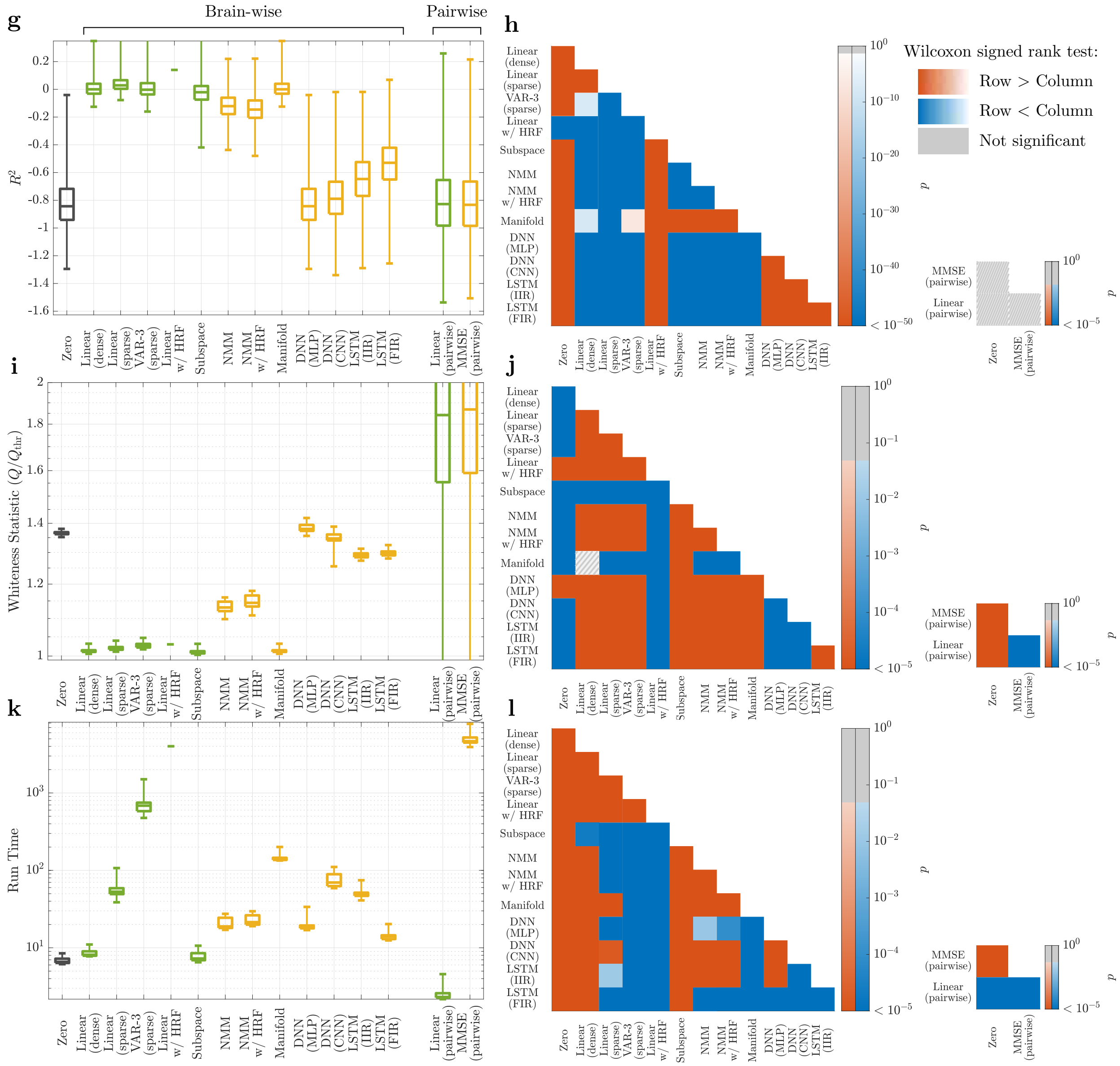


**Supplementary Figure 12: Linear vs. nonlinear models of unparcellated cortical rsfMRI activity.** To reduce computational complexity and be able to fit and validate all model families, we considered only vertices taken from two randomly selected cortical parcels: **(a-f)** a left somato-motor area '17Networks_LH_SomMotB_S2' consisting of 152 vertices, and **(g-l)** a right precuneus/posterior cingulate cortex area '17Networks_RH_DefaultA_pCunPCC_1' consisting of 177 vertices [5]). Panels and details parallel those in Supplementary Figure 11.

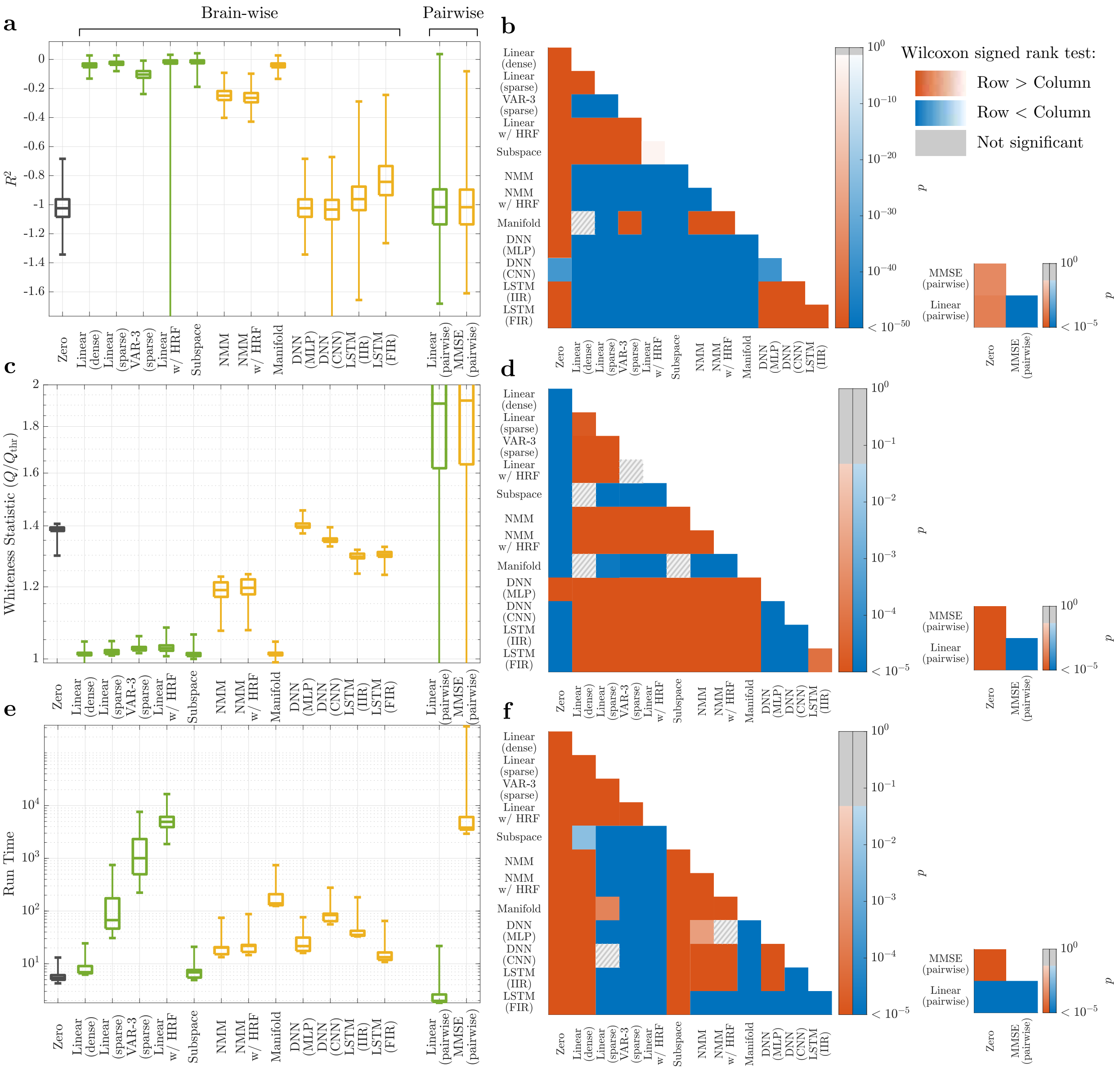


**Supplementary Figure 13: Linear vs. nonlinear models of unparcellated subcortical rsfMRI activity.** To reduce computational complexity and be able to fit and validate all model families, we considered only voxels from one randomly selected subcortical parcel (left dorsoanterior caudate, 'CAU-DA-lh' [6], consisting of 154 voxels). Panels and details parallel those in Supplementary Figure 11.

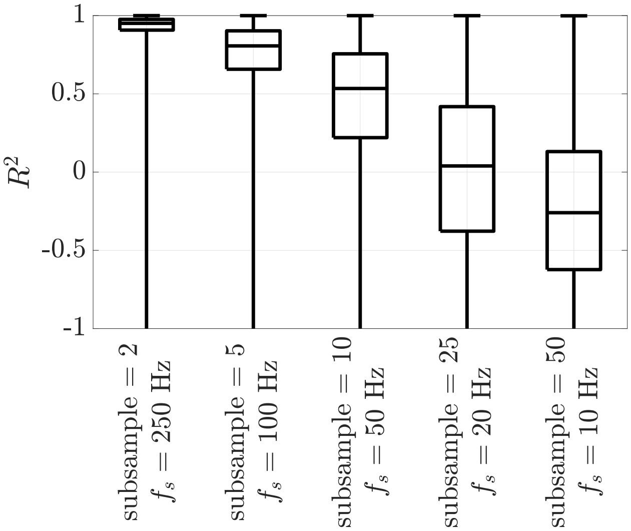


**Supplementary Figure 14: The channel-wise $R^2$ distribution of the zero model for iEEG data with different subsampling ratios and the corresponding sampling frequency.** As expected, higher subsampling results in less smooth time series, which in turn results in lower 'Zero' $R^2$, but also allows for using data from longer time intervals in model fitting and validation with the same amount of memory.

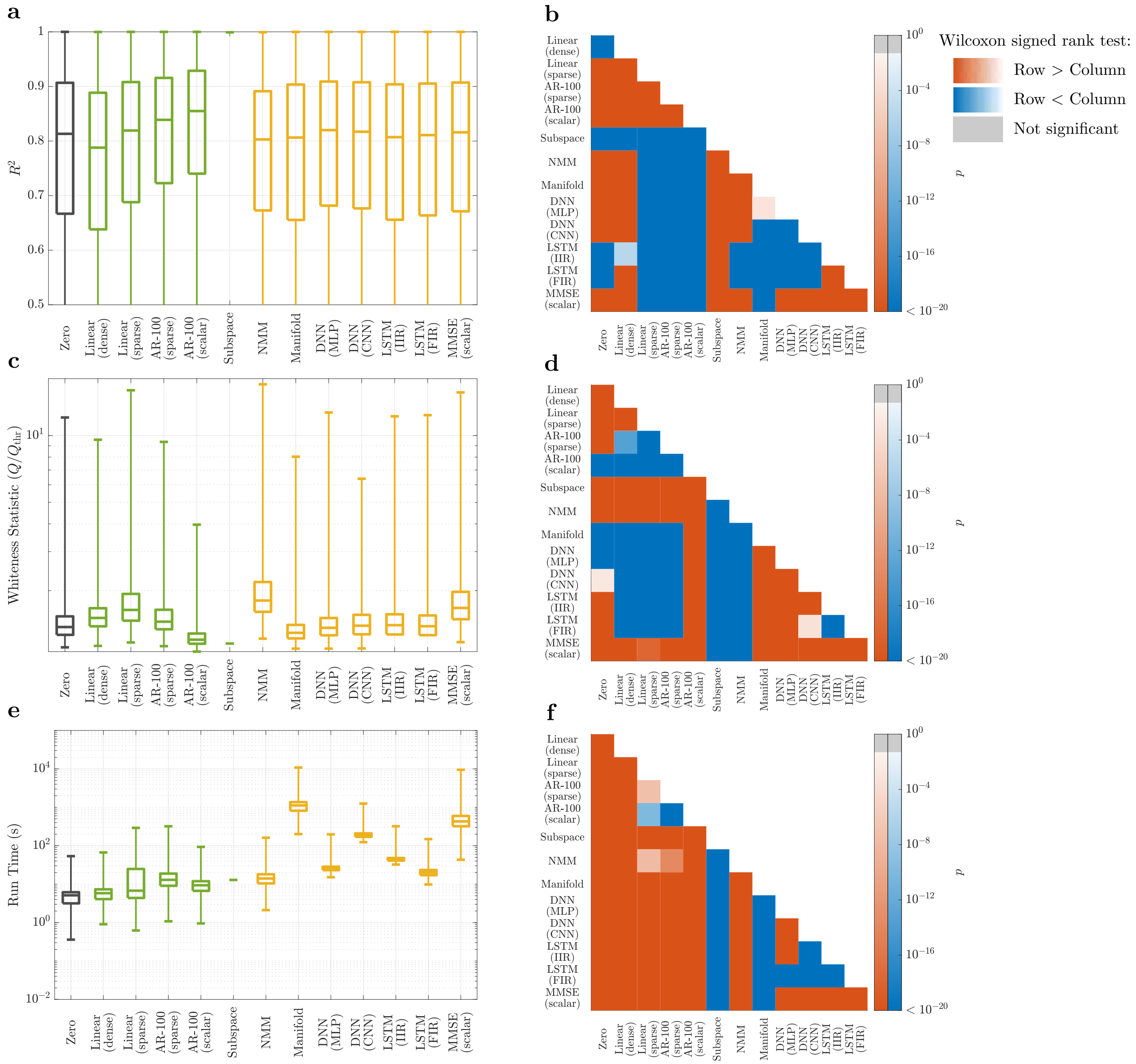

**Supplementary Figure 15: Linear versus nonlinear models of 5-fold subsampled rsiEEG activity.** Panels parallel those in Figure 3 in the main text. In panel (a), the box plot for the subspace method only has the top line (100th percentile) because for more than 75% of data segments the subspace method was unable to complete (either hung indefinitely or caused MATLAB to crash). For such cases we assign $R^2 = -\infty, Q = +\infty$, run time $= +\infty$ for the subspace method, causing its boxplot to miss the third quartile and anything below that. A similar situation holds in panels (c) and (e).

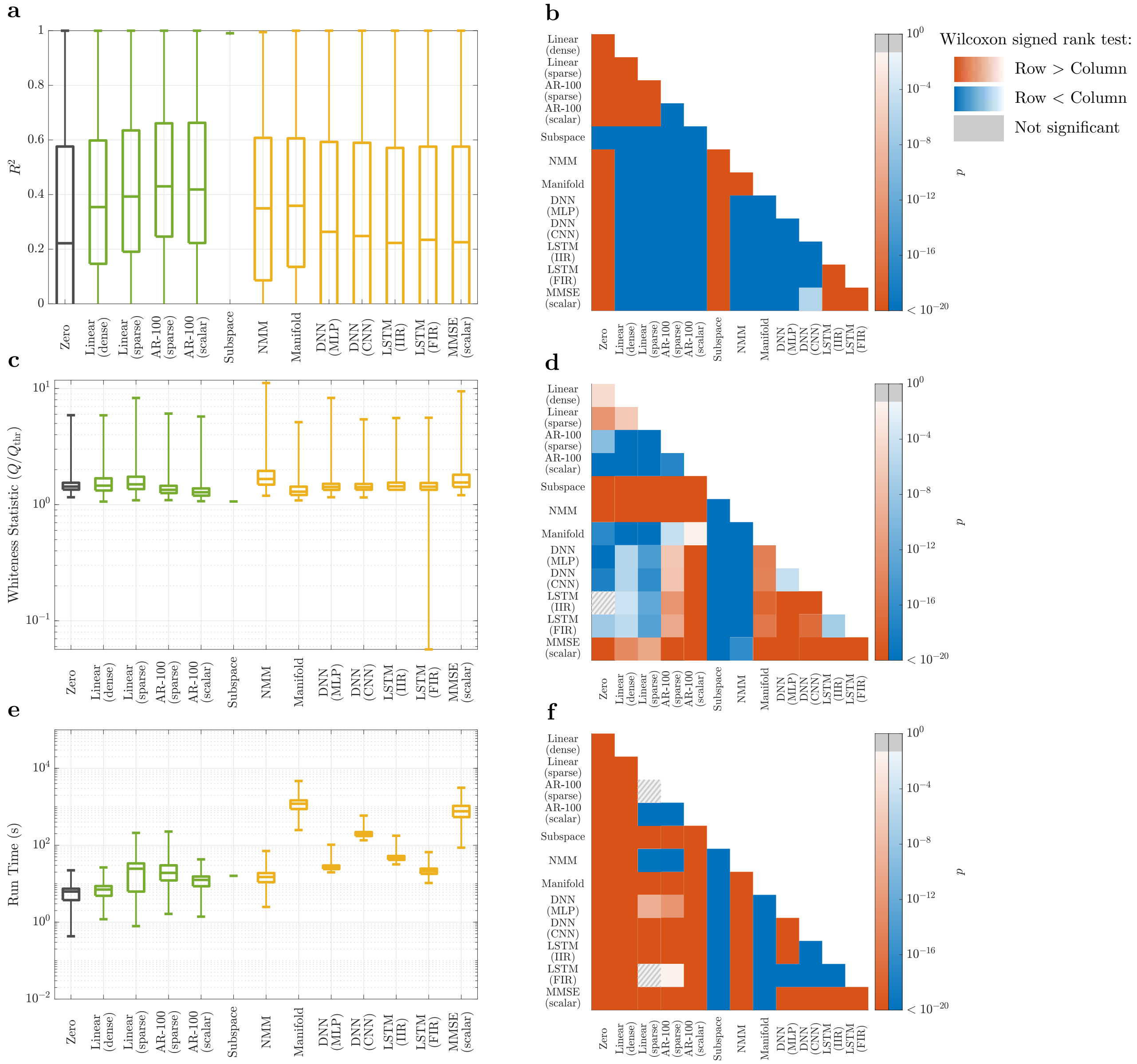


**Supplementary Figure 16: Linear versus nonlinear models of 25-fold subsampled rsiEEG activity.** Panels and details parallel those in Supplementary Figure 16. Note that here the 'AR-100 (sparse)' (which includes network interactions) has the highest $R^2$ distribution, even though the 'AR-100 (scalar)' model still has the whitest residuals.
